# Chemically modified guide RNAs enhance CRISPR-Cas13 knockdown in human cells

**DOI:** 10.1101/2021.05.12.443920

**Authors:** Alejandro Méndez-Mancilla, Hans-Hermann Wessels, Mateusz Legut, Anastasia Kadina, Megumu Mabuchi, John Walker, G. Brett Robb, Kevin Holden, Neville E. Sanjana

**Affiliations:** New York Genome Center, New York, NY, USA; Department of Biology, New York University, New York, NY, USA; Synthego Corporation, Redwood City, CA, USA; New England Biolabs, Ipswich, MA, USA

**Author notes:** These authors contributed equally (Alejandro Méndez-Mancilla, Hans-Hermann Wessels).

## Abstract

RNA-targeting CRISPR-Cas13 proteins have recently emerged as a powerful platform to transiently modulate gene expression outcomes. However, protein and CRISPR RNA (crRNA) delivery in human cells can be challenging and knockdown can be transient due to rapid crRNA degradation. Here we compare several chemical RNA modifications at different positions to identify synthetic crRNAs that improve RNA targeting efficiency and half-life in human cells. We show that co-delivery of modified crRNAs and recombinant Cas13 enzyme in ribonucleoprotein (RNP) complexes enables transient gene expression modulation in primary CD4+ and CD8+ T-cells. This system represents a robust and efficient method to transiently modulate transcripts without genetic manipulation.

---

The CRISPR-Cas13 enzyme family has shown remarkable versatility in basic biology and biotechnology applications^1^, including RNA knockdown^1–4^, transcript modifications like RNA editing^4,5^, live imaging^6,7^ and diagnostics^8–10^. Importantly, CRISPR-based transcriptome modulation has been proposed to offer considerable therapeutic potential for a wide spectrum of RNA-mediated diseases^1,11,12^. In all cases, a CRISPR RNA (crRNA) guides Cas13 to its target RNA by RNA-RNA hybridization of a short 23 nt spacer sequence to the target site^13,14^. Cas13-based effectors act on the RNA level and do not result in permanent changes to the genome. However, without a continuous source of crRNA expression, effects are short-lived due to rapid crRNA degradation by endogenous RNA nucleases and regeneration of the cellular steady-state by continuous target RNA expression.

Although continuous crRNA expression can be achieved via genetic manipulation, such as viral vectors expressing the crRNA and/or Cas13^2,3,15^, these methods are less desirable for more physiologically-relevant or therapeutic settings, such as targeting of immune checkpoints for immuno-oncology or differentiation of autologous stem cells prior to transplantation ^16,17^ . One of the main challenges for transcriptome manipulations is to achieve efficient delivery of CRISPR systems for robust RNA manipulation without modifying host DNA sequence. Recently, others have demonstrated chemical activation of crRNAs for Cas13a cleavage *in vitro*^18^ and delivered recombinant Cas13d into zebrafish embryos^19^. However, it remains unclear if a similar approach in human cells can lead to lasting effects. Inspired by guide RNA modifications for DNA-targeting Cas9 and Cas12a nucleases^20–26^, we sought to test if chemically modified synthetic crRNAs can be delivered into human cells to enhance Cas13-mediated transcript knockdown.

First, we assessed the degree of target RNA knockdown efficiency upon exogenous delivery of unmodified and chemically-modified crRNAs. We synthesized crRNAs with different chemical modifications at three uridine nucleotides (3xU) at the 3′ end of a 23 nt spacer sequence or with an inverted thymidine (invT) capping the 3′ end (**Figure 1a**). We tested 3 different modifications of the 3xU bases which have been reported before to improve RNA stability and evade secondary immune responses^20,22,27^: 2′-O-methylation (M), phosphorothioate linkage (S), and 2′-O-methylation and phosphorothioate linkage (MS). Although all of these modified crRNAs add extra bases at the 3’ end, we previously demonstrated that nucleotide mismatches to the target site beyond the 23 nt RNA-RNA hybridization interface do not interfere with target knockdown efficiency^14^. We chose to assess target knockdown efficiency of three broadly expressed cell surface proteins (CD46, CD55, CD71) that can be efficiently targeted with *Rfx*Cas13d^14^. The synthesized crRNAs were nucleofected into monoclonal *Rfx*Cas13d-expressing HEK293FT cells and, after three days, we quantified protein expression by flow cytometry (**Supplementary Figure 1a)**. For unmodified crRNAs, we noticed that protein knockdown for each of the three target transcripts was barely detectable relative to non-targeting crRNAs, suggesting that indeed unmodified crRNAs get rapidly cleared in human cells and cannot yield lasting knockdown effects (**Figure 1b, c, Supplementary Figure 1b**). On the other hand, all of the chemically modified crRNAs improved target knockdown but did so to varying degrees (one-way ANOVA, *p* < 10^−4^). We found that M-modified crRNAs had the overall largest knockdown (80% knockdown of CD71) but that there was greater variability among targeted transcripts. In contrast, we found that the invT modification improved knockdown efficiency in a more consistent manner across the three cell surface markers (55%) (**Figure 1c**). Surprisingly, the combination of M and S modifications did not result in improved knockdown than either individual (M or S) modification (one-way ANOVA, Tukey-corrected *p* = 0.14 [M], *p* = 0.89 [S]).

**Figure 1.**
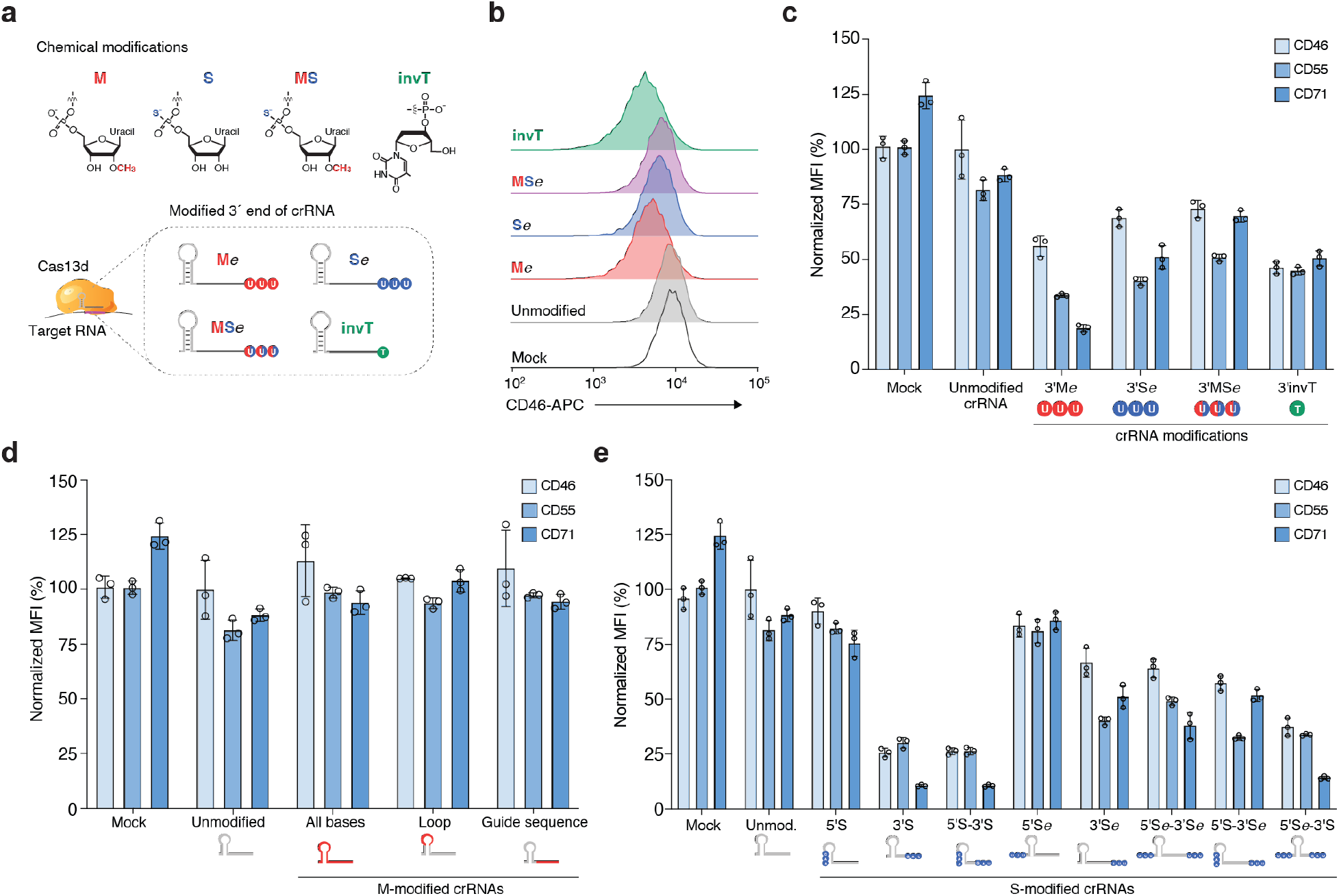
Chemically-modified CRISPR RNAs (crRNAs) improve Cas13 knockdown efficacy in human cells. **(a)** Overview of chemical modifications incorporated during synthesis of crRNAs: M, 3’-O-methyl base; S, phosphorothioate bond; MS, 3’-O-methyl base and phosphorothioate bond; invT, inverted thymidine. All crRNA sequences and modifications are listed in *Supplementary Table 1*. **(b)** Representative flow cytometry results for CD46 knockdown in HEK293FT-TetO-*Rfx*Cas13d-NLS cells nucleofected with synthetic crRNAs targeting CD46 — specifically, three modified uridines (M*e*, S*e*, MS*e* modifications with *e* denoting an extended crRNA with extra uridines) or an inverted thymidine directly following the last base of the guide sequence (invT modification). **(c, d, e)** Percent of CD46, CD55 and CD71 protein expression 72 hours post-nucleofection with the indicated crRNAs in HEK293FT-TetO-*Rfx*Cas13d-NLS cells. Cas13 expression was induced with doxycycline 24 hours prior to crRNA nucleofection. Target protein expression was measured by flow cytometry. RNA modification and placement were as follows: **(c)** synthetic crRNAs with chemical modifications located in the 3’ crRNA end — specifically, three modified uridines (M, S, MS modifications) or directly following the last base of the guide sequence (invT modification); **(d)** synthetic crRNAs with 3’-O-methyl (M) modification in all nucleotides in the crRNA (direct repeat and guide), in 3 nt in the loop region of the direct repeat, or in the 23 nt guide sequence; and **(e)** synthetic crRNAs chemically modified with a phosphorothioate bond placed in the crRNA sequence at the 5’ and/or 3’ end (S), or in three uridines added at the 5’ and/pr 3’ end (S*e*). Relative protein expression was calculated by comparing the normalized median fluorescent intensity (MFI) to cells nucleofected with non-targeting (NT) crRNAs. Bars represent mean values ± s.d., *n* = 3 biological replicate nucleofections.

In addition to improving Cas13-mediated target knockdown, we wondered if chemical modifications can close the gap between low and high efficiency guide sequences. Previously we used a high-throughput crRNA screen to identify optimal Cas13 guide sequence design parameters and developed a scoring metric to predict crRNA efficiency^14,28^. We chose one low-scoring guide sequence per target gene in addition to the high-scoring guide sequence used above (**Supplementary Figure 2a**) and found that chemically modified crRNAs do not generally improve the activity of low-scoring guide sequences (**Supplementary Figure 2b**). This suggests that chemical modification may improve crRNA stability but does not improve guide sequence efficiency. Stability of the crRNA is a key element of efficient Cas13 knockdown: Using M-modified crRNAs, we found that substantial knockdown was only possible when Cas13 was already expressed (**Supplementary Figure 2c**), similarly to previously reports with Cas9^20^. This suggests that Cas13 protein must be available for efficient ribonucleoprotein (RNP) complex formation after crRNAs are introduced into cells. We found similar results when we introduced Cas13 as messenger RNA at different timepoints relative to crRNA nucleofection (**Supplementary Figure 2d, e**).

Encouraged by these results, we next examined whether alternative placement of modified bases could further improve crRNA stability and transcript knockdown. Specifically, we tested if more extensively M-modified crRNAs could increase crRNA stability without interfering with target knockdown, as previously shown for DNA-targeting Cas9 sgRNA^24–26^. We synthesized crRNAs containing a M-modification along all the bases in the crRNA (53 modified bases), all bases in the spacer sequence (23 modified bases) or in the direct repeat loop sequence (3 modified bases). For all three targeted genes, these more extensive M-modifications abrogated Cas13 knockdown compared to unmodified crRNAs, presumably by disrupting the Cas13-crRNA interaction (**Figure 1d**). With Cas9 guide RNAs, multiple groups have found that moderate modification at the ends of the guide RNA tend to work best^22,23^, although one group reported an enhanced guide RNA with a majority of bases modified (∼70%) for targeting hepatocytes *in vivo*^24^. In contrast, our results show that a high degree of chemical modification yields less Cas13d knockdown.

Since partial modifications at the 3′ end boost knockdown, we decided to more systematically test the effect of partial modifications on the crRNA 5′ and 3′ ends. We tested crRNAs with S-modification at the first and last three nucleotides of the crRNA’s 5′ and 3′ ends (5S and 3S, respectively) or using a 3xU 5′ or 3′ extension (5′Se and 3′Se, respectively). We also tested all combinations of 5′ and 3′ end modifications. We found marked improvement of target knockdown for crRNAs with phosphorothioate modifications at the 3’ end of the spacer sequence (**Figure 1e**). Interestingly, 3′ phosphorothioate modifications improved knockdown efficiency to a greater degree when placed within the spacer sequence (3′S) compared to being placed in 3xU extensions (3′Se) (two-tailed *t*-test, *p* = 0.03). Modifications at the 5′ end of the crRNA alone, or in conjunction with 3’ modifications did not improve target knockdown efficiency (one-way ANOVA, Tukey-corrected *p* = 0.99 for all comparisons with added 5′ modifications). These results suggest that exonucleases or degradation processes at the crRNA 3′ end are a major hurdle for increased Cas13 activity using synthetic crRNAs^29^.

Next, we sought to assess the temporal dynamics of Cas13 activity with synthetic crRNAs by comparing knockdown of CD46 over time for 3′ invT, 3′S and unmodified crRNAs relative to non-targeting crRNAs (**Figure 2a**). All three targeting crRNAs, including unmodified crRNAs, yielded almost complete CD46 knockdown at 24 hours after nucleofection with around 95% protein loss for 3′S-modified crRNAs (**Figure 2b**). While CD46 expression quickly recovered in cells targeted with unmodified crRNAs, 3′S crRNAs led to pronounced knockdown at 48 hrs after crRNA delivery (**Figure 2c**). Even 4 days after nucleofection, the modified crRNAs resulted in ∼40% knockdown. Both crRNA modifications extend knockdown effects by about 2 days compared to unmodified crRNAs.

**Figure 2.**
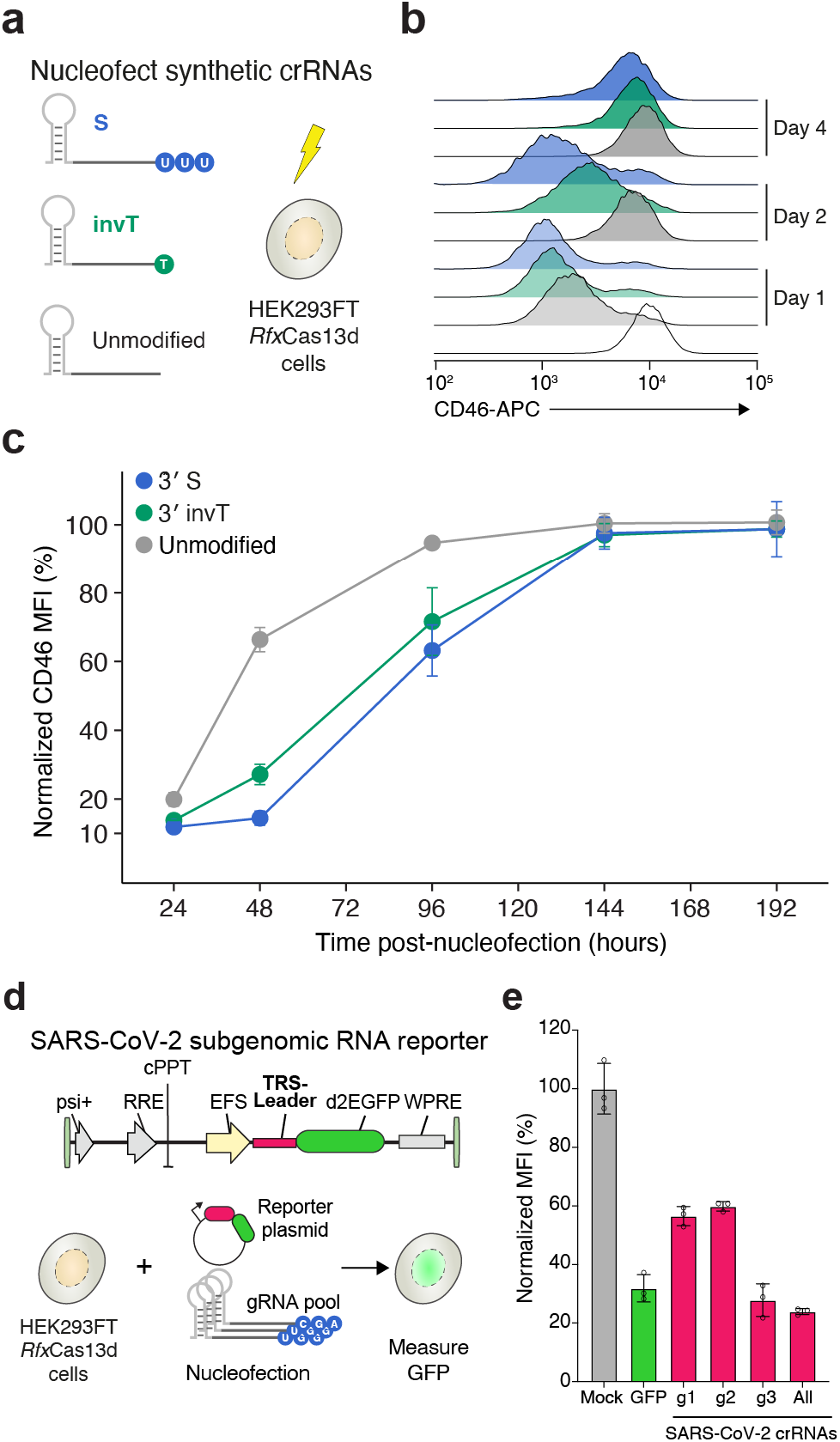
Transcript knockdown with chemically-modified crRNAs is sustained over multiple days. **(a)** Experimental design to measure the temporal dynamics of transient CD46 knockdown by nucleofecting the synthetic crRNAs in the HEK293FT-TetO-*Rfx*Cas13d-NLS cells. Synthetic crRNAs were unmodified, chemically modified with a phosphorothioate bond (S) on three uridines at the 3’ end, or chemically modified with an inverted thymidine at the 3’ end (invT). **(b)** Representative CD46 histograms at 1, 2 and 4 days after synthetic crRNA nucleofection. **(c)** Relative CD46 protein expression upon nucleofection with the synthetic crRNAs in panel *a*, normalized to cells nucleofected with non-targeting NT crRNAs. Points represent mean values ± s.d., *n* = 3 biological replicate nucleofections. **(d)** Experimental approach to measure SARS-CoV-2 RNA knockdown using a subgenomic RNA fluorescent reporter (destabilized EGFP) with synthetic crRNAs chemically modified with a phosphorothioate bond (S) on the three terminal nucleotides of the guide sequence. **(e)** EGFP expression in HEK293FT-TetO-*Rfx*Cas13d-NLS cells. Nucleofection conditions: *GFP*, pool of three S-modified crRNAs targeting EGFP; *g1, g2* and *g3*, individual S-modified crRNAs targeting the SARS-CoV-2 TRS-Leader sequence; *All*, pool of *g1, g2* and *g3* S-modified crRNAs. Percent GFP expression is determined relative to cells nucleofected with NT crRNAs. Bars represent mean values ± s.d., *n* = 3 biological replicate nucleofections.

Recently, several groups have suggested the possibility of using Cas13 to target severe acute respiratory syndrome coronavirus 2 (SARS-CoV-2), a positive-sense RNA virus responsible for the current global pandemic of novel coronavirus disease 2019 (COVID-19)^30–32^. To test if synthetic crRNA can degrade SARS-CoV-2, we designed a reporter construct that contains the SARS-CoV-2 leader sequence in the 5′ UTR of an EGFP reporter gene (**Figure 2d**). The SARS-CoV-2 leader sequence is a component of all SARS-CoV-2 subgenomic RNAs and thus represents a universal targeting site for all subgenomic viral transcripts ^33,34^. Similar to our experiments with endogenous human transcripts, we found that 3′S-modified crRNAs targeting the SARS-CoV-2 leader sequence can suppress reporter protein expression, despite targeting an untranslated sequence (**Figure 2e**). These results suggest that Cas13 together with chemically modified crRNAs may represent an efficient and programmable therapeutic approach to target universal SARS-CoV-2 sequences.

In addition to delivery of chemically-modified crRNAs for extended knockdown, another major challenge is delivery of the effector protein into systems that cannot easily be genetically modified to continuously express Cas13. Therefore, we next sought to evaluate if we could preassemble Cas13 RNP complexes with synthetic crRNAs and deliver these RNPs into primary human cells. First, we confirmed that preassembled RNPs with recombinant *Rfx*Cas13d with nuclear localization signals (NLSs) on the N- and C-termini (rCas13d) could efficiently degrade different target RNAs *in vitro* (**Figure 3a, Supplementary Figure 3a-c**). We found that purified rCas13d with or without an N-terminal MKIEE expression and solubility tag resulted in similar nuclease activity *in vitro*. By nucleofection of RNPs into human HEK293FT cells, we identified optimal buffer and temperature conditions for RNP assembly, achieving ∼85% knockdown of CD46 at 24 hours post-nucleofection (**Figure 3b, Supplementary Figure 3d**). At 48 hours post-nucleofection, we found that complexing in a Tris-based buffer at an elevated temperature (37°C) led to greater knockdown than an alternative buffer and room temperature RNP formation (two-way ANOVA with Tukey-corrected *p* < 10^−4^ for buffer and *p* < 10^−3^ for temperature). We further optimized RNP delivery using different protein amounts, while holding the molar ratio of rCas13d to crRNA (1:2) constant. We found that 10µg of rCas13d protein yielded the strongest knockdown and that additional rCas13d does not lead to greater knockdown (two-tailed *t*-test, *p* = 0.7) (**Figure 3c**). In agreement with our prior results using viral delivery of Cas13d, we find that using chemically-modified crRNAs also improves RNA knockdown with Cas13d RNPs (two-tailed paired *t*-test, *p* = 5 × 10^−3^).

**Figure 3.**
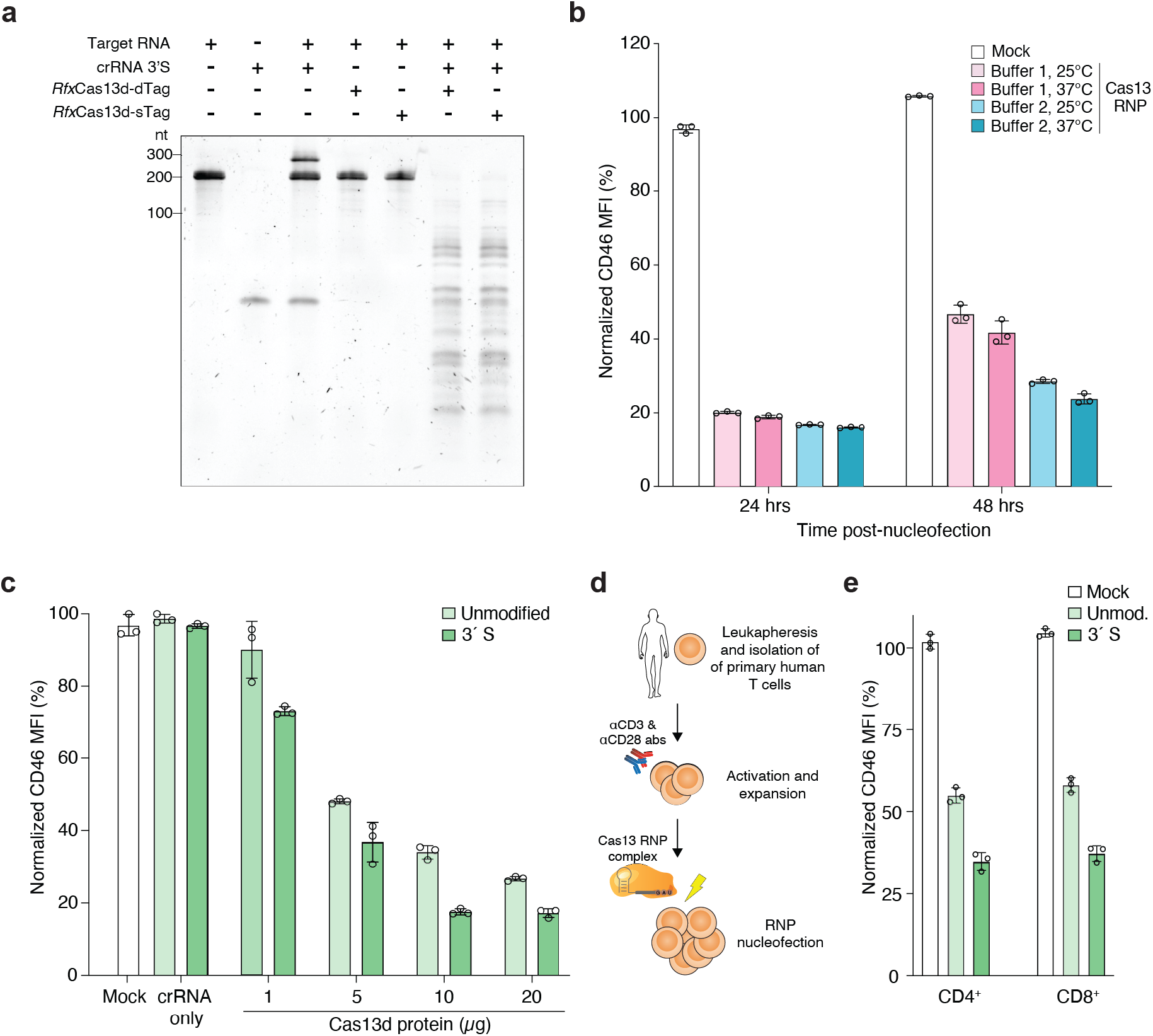
Robust knockdown in human cell lines and primary immune cells using Cas13d ribonucleoprotein (RNP) complexes and chemically-modified crRNAs. **(a)** Denaturing TBE-urea gel showing the cleavage activity of the indicated *Rfx*Cas13d proteins tagged with either one (sTag) or two (dTag) affinity purification tags and their cleavage activity using a chemically-modified crRNA (3’S) targeting CD46. *Rfx*Cas13d-sTag includes only a C-terminal HA tag. *Rfx*Cas13d-dTag includes C-terminal HA and 6xHis tags and an N-terminal MKIEE solubility sequence. **(b)** CD46 expression in HEK293FT cells at 24 and 48 hours after nucleofection with Cas13 RNP complexed with Buffer 1 (GenScript) and Buffer 2 buffer (RNA cleavage buffer, see *Methods*) at the indicated temperature prior to nucleofection. **(c)** CD46 expression in HEK293FT cells at 24 hours after nucleofection with synthetic crRNAs only and Cas13 RNP complexes with different protein amounts (1, 5, 10 and 20 µg) complexed with CD46-targeting synthetic crRNAs with no chemical modification or with a phosphorothioate bond at the 3′ end (3′S). **(d)** Leukapheresis and activation of primary human T cells prior to Cas13 RNP electroporation. **(e)** CD46 expression in primary human T cells (CD4+ and CD8+) at 24 hours after nucleofection with Cas13 RNPs complexed with CD46-targeting synthetic crRNAs with no chemical modification or with a 3′S modification. Bars represent mean values ± s.d., *n* =3 biological replicate nucleofections.

Using these optimized conditions for RNP formation and nucleofection, we isolated human T-cells from healthy donors, stimulated the cells with anti-CD3 and anti-CD28 antibodies, and asked whether we could knockdown CD46 expression in primary T-cells using these RNP complexes (**Figure 3d**). We assessed CD46 expression in CD4+ and CD8+ populations separately 24 hours after nucleofecting activated T cells (**Figure 3e, Supplementary Figure 3e**). For both T-cell populations we achieved a 60 – 65% knockdown of CD46 with 3′S modified crRNAs (**Figure 3f**). With unmodified crRNAs, we achieved only 40 – 45% knockdown of CD46. Taken together, these experiments demonstrate that Cas13 RNPs can modify gene expression in primary immune cells and that modified crRNAs lead to greater gene knockdown than unmodified crRNAs.

Here, we show that chemically-modified crRNAs can be used for Cas13d RNA targeting and knockdown. By testing different chemical modifications and their placement within the crRNA, we identify a subset of optimal modifications for RNA targeting in cells and demonstrate that knockdown effects persist longer than with unmodified crRNAs. Furthermore, we demonstrate the use of Cas13d RNPs complexes together with chemically-modified crRNAs in human cell lines and primary immune cells. Although chemically-modified crRNAs improve Cas13 knockdown with Cas13 from an integrated transgene or from RNPs, the improvement is more modest for RNPs (∼2-fold vs. 4-fold), which suggests that the RNP complex may also protect the crRNA from degradation. Given the recent development of Cas13-based research tools, diagnostics and therapeutics^2,19,30,35–37^, chemically-modified crRNAs can further enhance CRISPR-Cas13 RNA editing for diverse applications in biotechnology and medicine.

## Acknowledgements

We thank the entire Sanjana laboratory for support and advice. N.E.S. is supported by NYU and NYGC startup funds, NIH/NHGRI (DP2HG010099), NIH/NCI (R01CA218668), NIH/NIGMS (R01GM138635), DARPA (D18AP00053), the Cancer Research Institute, and the Brain and Behavior Foundation. M.L. is a Hope Funds for Cancer Research fellow.

## Authors contributions

A.M.-M, H.H.W and N.E.S conceived the project. A.M.-M., H.H.W., and N.E.S designed the experiments. A.M.-M. and H.H.W. performed *in vitro* and *in vivo* experiments. A.M.-M. and M.L. designed and performed experiments in primary T cells. A.K., J.W., and K.H. synthesized the crRNAs. M.M. and G.B.R. produced recombinant Cas13d protein. A.M.-M, H.H.W and N.E.S wrote the manuscript with input from all of the authors.

## Data and materials availability

Plasmids (TRS-Leader-SARS-CoV-2-d2eGFP, pET28-His-MBP-NLS-RfxCas13d-NLS-HA and pET28-MKIEE-NLS-RfxCas13d-NLS-HA-6His) have been deposited with Addgene (plasmid nos. 171585, 171586, 171587).

## Methods

### Plasmids

We cloned the TRS-Leader-SARS-CoV-2-d2eGFP plasmid (Addgene 171585) by introducing 75 nt of the SARS-CoV-2 Leader sequence into the *Nhe*I-digested EFS-EGFPd2PEST-2A-Hygro plasmid (Addgene 138152). We inserted the TRS-Leader immediately before the coding sequence by ligation of annealed oligos (see *Supplementary Table 3* for sequences). We generated bacterial *Rfx*Cas13d expression vectors as follows. Codon-optimized cDNAs containing the effector protein *Rfx*Cas13d were synthesized (Genscript) and assembled into modified pET28 vectors by NEBuilder HiFi DNA assembly (E2621, NEB). We created 2 bacterial expression vectors: pET28-His-MBP-NLS-RfxCas13d-NLS-HA (Addgene 171586) produces the protein *Rfx*Cas13d-sTag after cleavage of the N-terminal His-MBP fusion partner, and pET28-MKIEE-NLS-RfxCas13d-NLS-HA-6His (Addgene 171587) produces the protein *Rfx*Cas13d-dTag. For both constructs, NLS indicates the nuclear localization sequence derived from the simian virus 40 (SV40) Large T-antigen.

### Cell line culture conditions

HEK293FT cells were acquired from Thermo Fisher Scientific (R70007) and maintained in D10 media: DMEM with high glucose and stabilized L-glutamine (Caisson DML23) supplemented with 10% Serum Plus II fetal bovine serum (Sigma-Aldrich 14009C) and no antibiotics. For Tet-free D10 media, we omitted the Serum Plus II and instead substituted 10% tetracycline-negative fetal bovine serum (Corning 35-075-CV) for the Serum Plus II. HEK293FT were cultured at 37°C, 5% CO_2_, and ambient oxygen levels. We generated doxycycline-inducible HEK293FT-*Rfx*Cas13d-NLS cells using lentiviral transduction^17^.

### T cell isolation and culture conditions

T cells were procured from a de-identified healthy donor LeukoPak (New York Blood Center). Peripheral blood mononuclear cells (PBMCs) were isolated using Lymphoprep density gradient centrifugation (StemCell Technologies). CD8+ T cells were isolated from PBMCs by positive magnetic selection using EasySep Human CD8 Positive Selection Kit (StemCell Technologies). CD4+ T cells were isolated from PBMCs by negative magnetic selection using EasySep Human CD4 T cell Isolation Kit (StemCell Technologies). Isolated T cells were then plated in ImmunoCult-XF T Cell Expansion Medium (StemCell Technologies) supplemented with 10 ng μL-1 recombinant human IL-2 (StemCell Technologies) and activated with the ImmunoCult Human CD3/CD28 T Cell Activator (StemCell Technologies). T cells were cultured at 37°C, 5% CO_2_, and ambient oxygen levels.

### Chemically modified CRISPR RNA (crRNA) synthesis

The crRNAs were synthesized using solid-phase phosphoramidite chemistry (Synthego CRISPRevolution platform). Following synthesis, postprocessing and purification steps, we quantified each crRNA by UV absorption using Nanodrop (Thermo). We confirmed their identities and quality using an Agilent 1290 Infinity II high-performance liquid chromatography (HPLC) system coupled with an Agilent 6530B Quadrupole time of-flight mass spectrometry (Agilent Technologies) in negative ion polarity mode.

### Recombinant Cas13d production and purification

Recombinant *Rfx*Cas13d proteins were expressed in *E*.*coli* NiCo21(DE3) (NEB C2529). Cells were grown in 1L of lysogeny (Luria-Bertrani) broth containing 40 mg/mL kanamycin with shaking at 30°C. Protein expression was induced by the addition of 0.4 mM IPTG and the temperature was reduced to 16°C for 16 hours. Pelleted cells were disrupted by sonication. Proteins were purified from the lysate supernatant using HiTrapDEAE^FF^ Sepharose, HisTrap HP and HiTrap HeparinHP columns on an Akta Go instrument (Cytiva). To produce sTag *Rfx*Cas13d, the N-terminal His-MPB fusion partner was cleaved with Sumo protease and removed. Purified *Rfx*Cas13d proteins were stored in 20mM Tris-HCl, pH7.5, 500mM NaCl, 1mM EDTA, 1mM DTT and 50% glycerol (v/v). Protein concentration was determined by Bradford assay (Bio-Rad).

### Nucleofection of crRNAs and ribonucleoprotein (RNP) complexes

For nucleofection experiments in HEK293FT-*Rfx*Cas13d cells, we seeded three replicates of ∼2×10^6^ cells in D10 media with 1 ug/mL of doxycycline 24 hours before nucleofection. Using 1×10^5^ cells per condition/nucleofection, we nucleofected HEK293FT-*Rfx*Cas13d cells with 225 μmol of synthetic crRNAs in 20μl reaction of SF Cell Line Nucleofector Solution (Lonza V4XC-2032) using the Lonza Nucleofector 4D (program CM-130). Immediately after nucleofection, we plated cells in pre-equilibrated D10 media (equilibrated to 37° C and 5% CO_2_). For SARS-CoV-2 RNA knockdown, we nucleofected HEK293FT-*Rfx*Cas13d with 0.4 µg of TRS-Leader-SARS-CoV-2-d2eGFP plasmid together with 225 µmol of synthetic crRNAs and performed flow cytometry 24 hours after nucleofection.

For RNP delivery experiments, purified *Rfx*Cas13d protein and crRNAs were assembled at a 1:2 molar ratio (Cas13 protein : crRNA) under two different RNP complexing conditions: 1) 10 mins at 25°C or 37°C with nuclease reaction buffer (GenScript) and 2) 15 mins at 24°C or 37°C with RNA cleavage buffer (25mM Tris pH 7.5, 1mM DTT, 6mM MgCl_2_). After complexing, the *Rfx*Cas13d RNPs were nucleofected into HEK293FT cells or primary T cells using 10^5^ to 10^6^ cells in 20μl of nucleofection solutions specific for each cell type. For HEK293FT nucleofection, cells were nucleofected in 20μl reaction of SF Cell Line Nucleofector Solution (Lonza V4XC-2032) using the Lonza Nucleofector 4D (program CM-130). Immediately after nucleofection, HEK293FT cells were plated in pre-equilibrated D10 media (equilibrated to 37° C and 5% CO_2_) and remained there until flow cytometry analysis. For T cells nucleofection, two days after activation CD4+ and CD8+ T cells were combined, washed 2X in Dulbecco’s PBS without calcium or magnesium (D-PBS, Caisson Labs) and resuspended in 20 μl P3 Primary Cell Nucleofector Solution (Lonza V4XP-3032). T cells were then nucleofected using the Lonza Nucleofector 4D (program E0-115). Nucleofections were performed in triplicate. Immediately after nucleofection T cells were plated in pre-equilibrated ImmunoCult-XF T Cell Expansion Medium supplemented with 10 ng μL-1 recombinant human IL-2 (equilibrated to 37° C and 5% CO_2_). Before flow cytometry analysis, nucleofected T cells were washed with D-PBS and stained with LIVE/DEAD Fixable Violet Dead Cell Stain (ThermoFisher) for 5 minutes at room temperature in the dark before proceeding with antibody stain.

### Biochemical in vitro RNA cleavage assays

We synthesized 200 bp target RNAs for CD46, CD55 and CD71 by PCR amplification of templates from total cDNA from HEK293FT cells and then performed T7 *in vitro* RNA transcription. To prepare the cDNA library, we extracted total RNA from HEK293FT cells using Direct-zol purification (Zymo) and reverse transcribed 1 ug of RNA into cDNA using Revert-Aid (ThermoFisher). Using the PCR templates of target RNAs, we performed *in vitro* RNA transcription using the HiScribe T7 High Yield RNA Synthesis Kit (NEB). After transcription, RNA was purified using a Monarch RNA Cleanup Kit (NEB) and quantified on a Nanodrop spectrophotometer (Thermo).

Similarly, *Rfx*Cas13d and eGFP messenger RNAs (mRNAs) were synthesized using a T7 RNA polymerase *in vitro* transcription with the HiScribe T7 ARCA mRNA Kit (with tailing) (NEB). The mRNAs were purified using a Monarch RNA Cleanup Kit (NEB). We amplified target RNAs and mRNAs using primers that include a T7 promoter sequence on the 5′ end; for the mRNAs, we also included the corresponding Kozak and start/stop codon sequences (**Supplementary Table 2)**.

Purified *Rfx*Cas13d proteins and synthetic crRNAs were mixed (unless otherwise indicated) at 2:1 molar ratio in Buffer 1 (GenScript SC1841) or Buffer 2^2^ (25mM Tris pH 7.5, 1mM DTT, 6mM MgCl_2_). The reaction was prepared on ice and incubated at 25° C or 37° C for 15 minutes prior to the addition of target RNA at 1:2 molar ratio relative to *Rfx*Cas13d protein. The reaction was incubated at 37° C for 45 minutes and quenched with 1 μL of enzyme stop solution^2^ (10 mg/mL Proteinase K, 4M Urea, 80mM EDTA, 20mM Tris pH 8.0) at 37° C for 15 minutes. The samples were denatured at 95°C for 5 minutes in 2X RNA Gel Loading Dye (ThermoFisher) and loaded onto 10% TBE-Urea gels (ThermoFisher). The gels were run at 180 V for 35 mins. After separation, gels were stained with SYBR Gold (ThermoFisher) prior to imaging via Gel Doc EZ system (Bio-Rad).

### Flow cytometry

Cells were stained with the following antibodies: PE anti-human CD4 (clone RPA-T4, Biolegend 300507), FITC anti-human CD8a (clone RPA-T8, Biolegend 301006) and APC anti-human CD46 (clone TRA-2-10, Biolegend 352405). Antibody staining was performed for 20 minutes on ice. After staining the cells were washed 2X in D-PBS and analyzed via flow cytometry (Sony SH800S). A minimum of 10,000 viable events was collected per sample.

### Data analysis

Data analysis and statistical testing was performed using GraphPad Prism 8 (GraphPad Software Inc.). Flow cytometry analysis and figure generation was performed using FlowJo v10 (BD). Specific statistical analysis methods are described where results are presented.

