## Supplementary Figures and Tables for "Chemically modified guide RNAs enhance CRISPR-Cas13 knockdown in human cells"

† These authors contributed equally.

**Supplementary figures and tables**

**Supplementary Figure 1.** Flow cytometry for Cas13 synthetic CRISPR RNA (crRNA) knockdown in mammalian cells.

**Supplementary Figure 2.** Effect of guide sequence prediction and timing of Cas13 induction on knockdown.

**Supplementary Figure 3.** Characterization of Cas13 ribonucleoproteins (RNPs) *in vitro* and *in vivo*.

**Supplementary Table 1.** Chemically-modified Cas13 CRISPR RNAs (crRNAs).

**Supplementary Table 2.** PCR primers for T7 *in vitro* transcription.

**Supplementary Table 3.** Primers for SARS-CoV-2 TRS-Leader sequence cloning.

**Supplementary Figure 1**

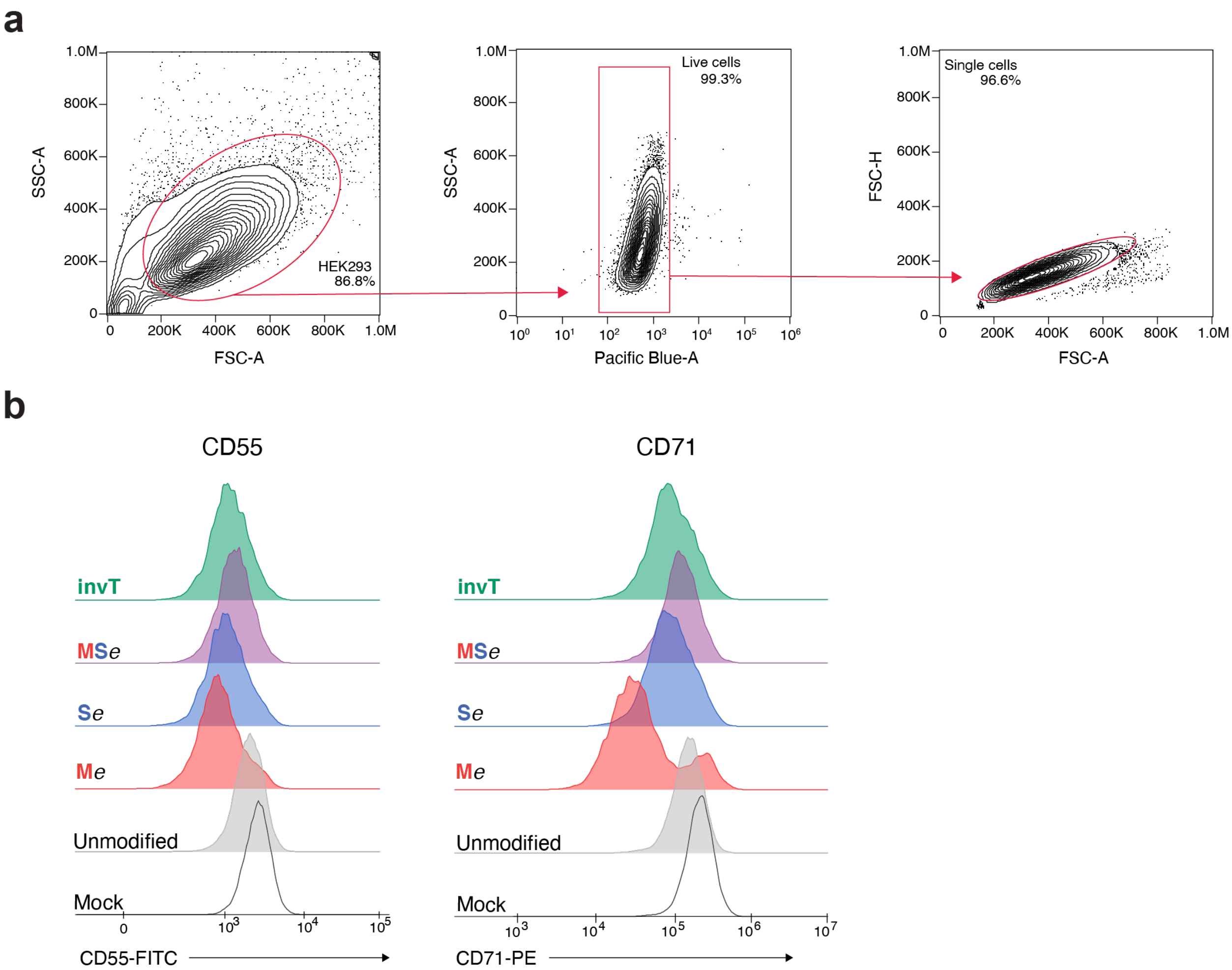


**Supplementary Figure 1. Flow cytometry for Cas13 synthetic CRISPR RNA (crRNA) knockdown in mammalian cells.**

(**a**) Flow cytometry gating strategy for CD46, CD55 and CD71 knockdown in HEK293FT-Tet-NLS-*Rfx*Cas13d-NLS cells nucleofected with synthetic crRNAs. (**b**) Representative histograms of CD55 and CD71 expression in HEK293FT-Tet-*Rfx*Cas13d-NLS cells nucleofected with the indicated synthetic crRNAs — specifically, three modified uridines (M*e*, S*e*, MS*e* modifications with *e* denoting an extended crRNA with extra uridines) or an inverted thymidine directly following the last base of the guide sequence (invT modification). M, 3'-O-methyl base; S, phosphorothioate bond; MS, 3'-O-methyl base and phosphorothioate bond; invT, inverted thymidine.

**Supplementary Figure 2**

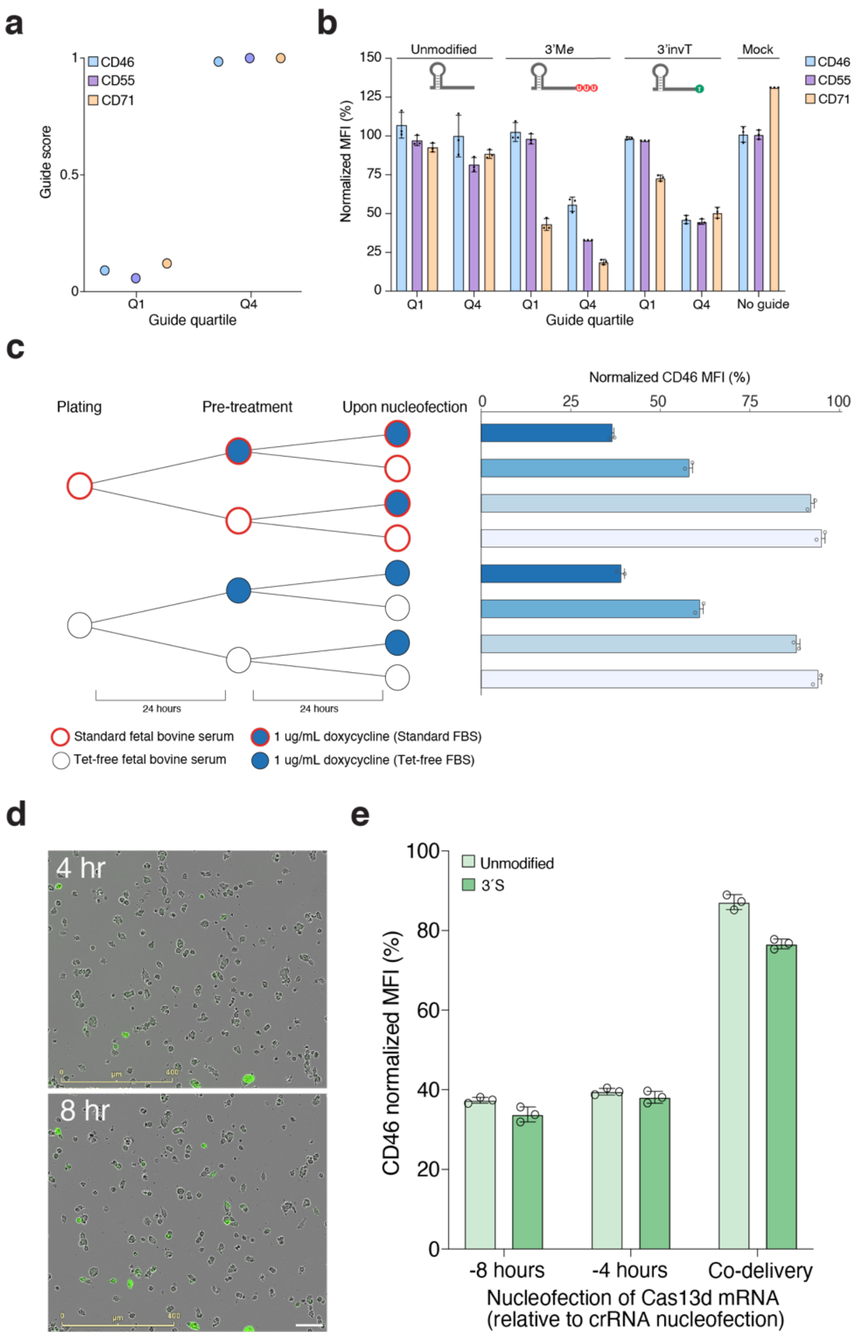


**Supplementary Figure 2. Effect of guide sequence prediction and timing of Cas13 induction on knockdown.**

(**a**) Predicted Cas13 guide RNA scores targeting CD46, CD55 and CD71. For each target, we used the *cas13design* webtool[14,28] to design 2 guide RNAs — one predicted as low activity (Q1) and one predicted as high activity (Q4). (**b**) Expression of CD46, CD55 and CD71 in HEK293FT-Tet-*Rfx*Cas13d-NLS cells nucleofected with the indicated synthetic crRNAs. Bars indicate the mean and error bars denote s.d., *n* = 3 nucleofection replicates. (**c**) CD46 knockdown in HEK293FT-Tet-NLS-*Rfx*Cas13d-NLS cells with different media and timing of doxycycline induction. Cells were grown in standard D10 media (*top*) or Tet-free D10 media (*bottom*) and doxycycline was added either 1 day prior to and/or upon crRNA nucleofection. Bars indicate the mean and error bars denote s.d., *n* = 2 nucleofection replicates. (**d**) Time-course of EGFP expression (mRNA transfection) in HEK293FT cells visualized by live imaging (Incucyte). $5.0 \times 10^5$ HEK293FT cells were nucleofected with 7.5µg of EGFP mRNA and then imaged at the times indicated post-transfection. The scale bar (lower-right white bar) in the bottom panel denotes 100 µm. (**e**) Staggered delivery of 7.5µg *Rfx*Cas13d mRNA and 1.125 mmol of CD46 synthetic crRNAs into $5.0 \times 10^5$ HEK293FT cells. The Cas13 mRNA was delivered 8 hours before, 4 hours before or simultaneously with the crRNA. At 24 hours post-nucleofection of the crRNA, we computed the normalized median fluorescent intensity (MFI) of CD46, relative to cells nucleofected with non-targeting crRNAs. Bars represent mean values ± s.d., *n* = 3 biological replicate nucleofections.

### Supplementary Figure 3

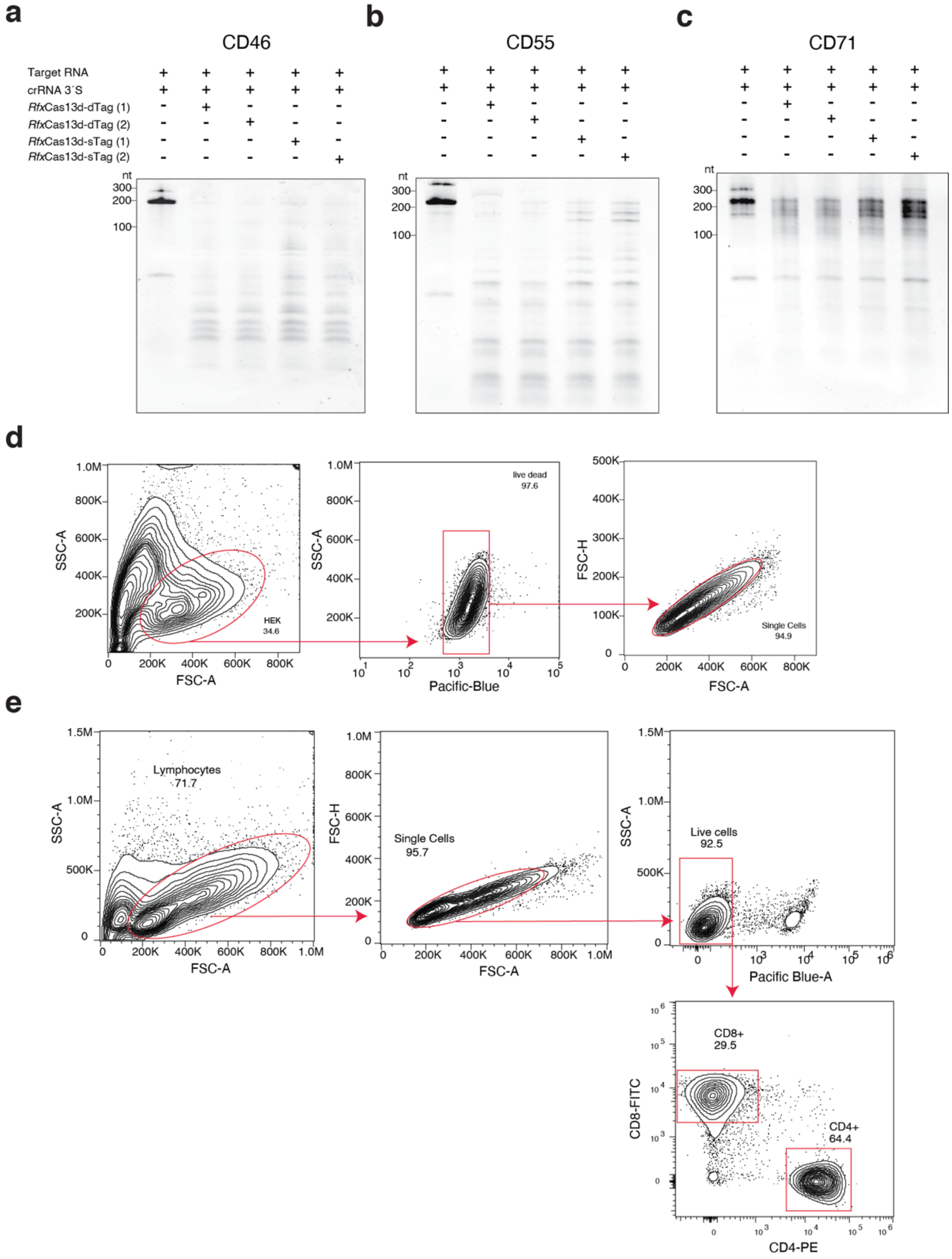


**Supplementary Figure 3. Characterization of Cas13 ribonucleoproteins (RNPs) *in vitro* and *in vivo*.**

**(a)** Denaturing RNA gel showing the cleavage activity of recombinant *Rfx*Cas13d proteins tagged with C-terminal HA tag (single-tag, or sTag), or with N-terminal MKIEE and C-terminal HA and 6xHis tags (double-tag, or dTag) and their cleavage activity using a chemically-modified crRNA

(3´S) targeting CD46. Two independent purifications are shown for each enzyme. **(b)** Denaturing RNA gel showing recombinant *Rfx*Cas13d proteins tagged with C-terminal HA tag (single-tag, or sTag) or with C-terminal HA tag (single-tag, or sTag), or with N-terminal MKIEE, C-terminal HA and 6xHis tags (double-tag, or dTag) and their cleavage activity using a chemically-modified crRNA (3´S) targeting CD46. Two independent purifications are shown for each enzyme. **(c)** Denaturing RNA gel showing recombinant *Rfx*Cas13d proteins tagged with C-terminal HA tag (single-tag, or sTag) or with N-terminal MKIEE, C-terminal HA and 6xHis tags (double-tag, or dTag) and their cleavage activity using a chemically-modified crRNA (3´S) targeting CD46. Two independent purifications are shown for each enzyme. **(d)** Flow cytometry gating strategy for CD46 knockdown using *Rfx*Cas13d-dTag RNPs with synthetic crRNAs in HEK293FT cells. **(e)** Flow cytometry gating strategy for CD46 knockdown using *Rfx*Cas13d-sTag RNPs with synthetic crRNAs in human primary CD4+ and CD8+ T cells.

### Supplementary Table 1. Chemically modified Cas13 crRNAs.

| No. | crRNA name | crRNA sequence |
|---|---|---|
| 1 | UnmodCD46 | AACCCCUACCAACUGGUCGGGGUUUGAAACAGACAAUUGUGUCGCUGCCAUCG |
| 2 | UnmodCD46 | AACCCCUACCAACUGGUCGGGGUUUGAAACGCGGCGGCCGGGAGGCUCCAUGC |
| 3 | 3'MeCD46 | AACCCCUACCAACUGGUCGGGGUUUGAAACAGACAAUUGUGUCGCUGCCAUCGmUmUmU |
| 4 | 3'MeCD46 | AACCCCUACCAACUGGUCGGGGUUUGAAACGCGGCGGCCGGGAGGCUCCAUGCmUmUmU |
| 5 | 3'SeCD46 | AACCCCUACCAACUGGUCGGGGUUUGAAACAGACAAUUGUGUCGCUGCCAUCGU*U*U |
| 6 | 3'SeCD46 | AACCCCUACCAACUGGUCGGGGUUUGAAACGCGGCGGCCGGGAGGCUCCAUGCU*U*U |
| 7 | 3'MSeCD46 | AACCCCUACCAACUGGUCGGGGUUUGAAACAGACAAUUGUGUCGCUGCCAUCGmU*mU*mU |
| 8 | 3'MSeCD46 | AACCCCUACCAACUGGUCGGGGUUUGAAACGCGGCGGCCGGGAGGCUCCAUGCmU*mU*mU |
| 9 | iCD46 | AACCCCUACCAACUGGUCGGGGUUUGAAACAGACAAUUGUGUCGCUGCCAUCGdT |
| 10 | iCD46 | AACCCCUACCAACUGGUCGGGGUUUGAAACGCGGCGGCCGGGAGGCUCCAUGCdT |
| 11 | FullMCD46 | mAmAmCmCmCmCmUmAmCmCmAmAmCmUmGmGmUmCmGmGmGmGmUmUmUmGmAmAmAmCmAmG mAmCmAmAmUmUmGmUmGmUmCmGmCmUmGmCmCmAmUmCmG |
| 12 | FullMCD46 | mAmAmCmCmCmCmUmAmCmCmAmAmCmUmGmGmUmCmGmGmGmGmUmUmUmGmAmAmAmCmGmC mGmGmCmGmGmCmCmGmGmGmAmGmGmCmUmCmCmAmUmGmC |
| 13 | GuideMCD46 | AACCCCUACCAACUGGUCGGGGUUUGAAACmAmGmAmCmAmAmUmUmGmUmGmUmCmGmCmUmG mCmCmAmUmCmG |
| 14 | GuideMCD46 | AACCCCUACCAACUGGUCGGGGUUUGAAACmGmCmGmGmCmGmGmCmCmGmGmGmAmGmGmCmU mCmCmAmUmGmC |
| 15 | UnmodCD55 | AACCCCUACCAACUGGUCGGGGUUUGAAACUCCCCGAGGAGGGGCAGCGCCGC |
| 16 | 3'MeCD55 | AACCCCUACCAACUGGUCGGGGUUUGAAACUCCCCGAGGAGGGGCAGCGCCGCmUmUmU |
| 17 | 3'SeCD55 | AACCCCUACCAACUGGUCGGGGUUUGAAACUCCCCGAGGAGGGGCAGCGCCGCU*U*U |
| 18 | 3'MSeCD55 | AACCCCUACCAACUGGUCGGGGUUUGAAACUCCCCGAGGAGGGGCAGCGCCGCmU*mU*mU |
| 19 | iCD55 | AACCCCUACCAACUGGUCGGGGUUUGAAACUCCCCGAGGAGGGGCAGCGCCGCdT |
| 20 | FullMCD55 | mAmAmCmCmCmCmUmAmCmCmAmAmCmUmGmGmUmCmGmGmGmGmUmUmUmGmAmAmAmCmUmC mCmCmCmGmAmGmGmAmGmGmGmGmCmAmGmCmGmCmCmGmC |
| 21 | GuideMCD55 | AACCCCUACCAACUGGUCGGGGUUUGAAACmUmCmCmCmCmGmAmGmGmAmGmGmGmGmCmAmG mCmGmCmCmGmC |
| 22 | UnmodCD71 | AACCCCUACCAACUGGUCGGGGUUUGAAACCGAGCCAGGCUGAACCGGGUAUA |
| 23 | 3'MeCD71 | AACCCCUACCAACUGGUCGGGGUUUGAAACCGAGCCAGGCUGAACCGGGUAUAmUmUmU |
| 24 | 3'SeCD71 | AACCCCUACCAACUGGUCGGGGUUUGAAACCGAGCCAGGCUGAACCGGGUAUAU*U*U |
| 25 | 3'MSeCD71 | AACCCCUACCAACUGGUCGGGGUUUGAAACCGAGCCAGGCUGAACCGGGUAUAmU*mU*mU |
| 26 | iCD71 | AACCCCUACCAACUGGUCGGGGUUUGAAACCGAGCCAGGCUGAACCGGGUAUAdT |
| 27 | FullMCD71 | mAmAmCmCmCmCmUmAmCmCmAmAmCmUmGmGmUmCmGmGmGmGmUmUmUmGmAmAmAmCmCmG mAmGmCmCmAmGmGmCmUmGmAmAmCmCmGmGmGmUmAmUmA |

| | | |
|---|---|---|
| 28 | GuideMCD71 | AACCCCUACCAACUGGUCGGGGUUUGAAACmCmGmAmGmCmCmAmGmGmCmUmGmAmAmCmCmG mGmGmUmAmUmA |
| 29 | UnmodNT | AACCCCUACCAACUGGUCGGGGUUUGAAACUACUUACCUAAUGCGCGGUAGAU |
| 30 | 3'MeNT | AACCCCUACCAACUGGUCGGGGUUUGAAACUACUUACCUAAUGCGCGGUAGAUmUmUmU |
| 31 | 3'SeNT | AACCCCUACCAACUGGUCGGGGUUUGAAACUACUUACCUAAUGCGCGGUAGAUU*U*U |
| 32 | 3'MSeNT | AACCCCUACCAACUGGUCGGGGUUUGAAACUACUUACCUAAUGCGCGGUAGAUmU*mU*mU |
| 33 | iNT | AACCCCUACCAACUGGUCGGGGUUUGAAACUACUUACCUAAUGCGCGGUAGAUdT |
| 34 | FullMNT | mAmAmCmCmCmCmUmAmCmCmAmAmCmUmGmGmUmCmGmGmGmGmUmUmUmGmAmAmAmCmUmA mCmUmUmAmCmCmUmAmAmUmGmCmGmCmGmGmUmAmGmAmU |
| 35 | GuideMNT | AACCCCUACCAACUGGUCGGGGUUUGAAACmUmAmCmUmUmAmCmCmUmAmAmUmGmCmGmCmG mGmUmAmGmAmU |
| 36 | 5'SCD46 | A*A*CCCCUACCAACUGGUCGGGGUUUGAAACGCGGCGGCCGGGAGGCUCCAUGC |
| 37 | 3'SCD46 | AACCCCUACCAACUGGUCGGGGUUUGAAACGCGGCGGCCGGGAGGCUCCAU*G*C |
| 38 | 5'-3'-SCD46 | A*A*CCCCUACCAACUGGUCGGGGUUUGAAACGCGGCGGCCGGGAGGCUCCAU*G*C |
| 39 | 5'SeCD46 | U*U*UAACCCCUACCAACUGGUCGGGGUUUGAAACGCGGCGGCCGGGAGGCUCCAUGC |
| 40 | 5'-3'-SeCD46 | U*U*UAACCCCUACCAACUGGUCGGGGUUUGAAACGCGGCGGCCGGGAGGCUCCAUGCU*U*U |
| 41 | 5'S-3'SeCD46 | A*A*CCCCUACCAACUGGUCGGGGUUUGAAACGCGGCGGCCGGGAGGCUCCAUGCU*U*U |
| 42 | 5'Se-3'SCD46 | U*U*UAACCCCUACCAACUGGUCGGGGUUUGAAACGCGGCGGCCGGGAGGCUCCAU*G*C |
| 43 | 5'SCD46 | A*A*CCCCUACCAACUGGUCGGGGUUUGAAACAGACAAUUGUGUCGCUGCCAUCG |
| 44 | 3'SCD46 | AACCCCUACCAACUGGUCGGGGUUUGAAACAGACAAUUGUGUCGCUGCCAU*C*G |
| 45 | 5'-3'-SCD46 | A*A*CCCCUACCAACUGGUCGGGGUUUGAAACAGACAAUUGUGUCGCUGCCAU*C*G |
| 46 | 5'SeCD46 | U*U*UAACCCCUACCAACUGGUCGGGGUUUGAAACAGACAAUUGUGUCGCUGCCAUCG |
| 47 | 5'-3'-SeCD46 | U*U*UAACCCCUACCAACUGGUCGGGGUUUGAAACAGACAAUUGUGUCGCUGCCAUCGU*U*U |
| 48 | 5'S-3'SeCD46 | A*A*CCCCUACCAACUGGUCGGGGUUUGAAACAGACAAUUGUGUCGCUGCCAUCGU*U*U |
| 49 | 5'Se-3'SCD46 | U*U*UAACCCCUACCAACUGGUCGGGGUUUGAAACAGACAAUUGUGUCGCUGCCAU*C*G |
| 50 | 5'SNT | A*C*CCCUACCAACUGGUCGGGGUUUGAAACUACUUACCUAAUGCGCGGUAGAU |
| 51 | 3'SNT | AACCCCUACCAACUGGUCGGGGUUUGAAACUACUUACCUAAUGCGCGGUAG*A*U |
| 52 | 5'-3'-SNT | A*A*CCCCUACCAACUGGUCGGGGUUUGAAACUACUUACCUAAUGCGCGGUAG*A*U |
| 53 | 5'SeNT | U*U*UAACCCCUACCAACUGGUCGGGGUUUGAAACUACUUACCUAAUGCGCGGUAGAU |
| 54 | 5'-3'-SeNT | U*U*UAACCCCUACCAACUGGUCGGGGUUUGAAACUACUUACCUAAUGCGCGGUAGAUU*U*U |
| 55 | 5'S-3'SeNT | A*A*CCCCUACCAACUGGUCGGGGUUUGAAACUACUUACCUAAUGCGCGGUAGAUU*U*U |
| 56 | 5'Se-3'SNT | U*U*UAACCCCUACCAACUGGUCGGGGUUUGAAACUACUUACCUAAUGCGCGGUAG*A*U |
| 57 | 3'MSeCD55 | AACCCCUACCAACUGGUCGGGGUUUGAAACGAUCACUGAGUCCUUCUCGCCAGmU*mU*mU |

| | | |
|---|---|---|
| 58 | 3'MSeCD71 | AACCCCUACCAACUGGUCGGGGUUUGAAACCGAUCACAGCAAUAGUCCCAUAGmU*mU*mU |
| 59 | UnmodCD55 | AACCCCUACCAACUGGUCGGGGUUUGAAACGAUCACUGAGUCCUUCUCGCCAG |
| 60 | 3'MeCD55 | AACCCCUACCAACUGGUCGGGGUUUGAAACGAUCACUGAGUCCUUCUCGCCAGmUmUmU |
| 61 | 3'SeCD55 | AACCCCUACCAACUGGUCGGGGUUUGAAACGAUCACUGAGUCCUUCUCGCCAGU*U*U |
| 62 | iCD55 | AACCCCUACCAACUGGUCGGGGUUUGAAACGAUCACUGAGUCCUUCUCGCCAGTd |
| 63 | FullMCD55 | mAmAmCmCmCmCmUmAmCmCmAmAmCmUmGmGmUmCmGmGmGmGmUmUmUmGmAmAmAmCmGmA mUmCmAmCmUmGmAmGmUmCmCmUmUmCmUmCmGmCmCmAmG |
| 64 | GuideMCD55 | AACCCCUACCAACUGGUCGGGGUUUGAAACmGmAmUmCmAmCmUmGmAmGmUmCmCmUmUmCmU mCmGmCmCmAmG |
| 65 | UnmodCD71 | AACCCCUACCAACUGGUCGGGGUUUGAAACCGAUCACAGCAAUAGUCCCAUAG |
| 66 | 3'MeCD71 | AACCCCUACCAACUGGUCGGGGUUUGAAACCGAUCACAGCAAUAGUCCCAUAGmUmUmU |
| 67 | 3'SeCD71 | AACCCCUACCAACUGGUCGGGGUUUGAAACCGAUCACAGCAAUAGUCCCAUAGU*U*U |
| 68 | iCD71 | AACCCCUACCAACUGGUCGGGGUUUGAAACCGAUCACAGCAAUAGUCCCAUAGTd |
| 69 | FullMCD71 | mAmAmCmCmCmCmUmAmCmCmAmAmCmUmGmGmUmCmGmGmGmGmUmUmUmGmAmAmAmCmCmG mAmUmCmAmCmAmGmCmAmAmUmAmGmUmCmCmCmAmUmAmG |
| 70 | GuideMCD71 | AACCCCUACCAACUGGUCGGGGUUUGAAACmCmGmAmUmCmAmCmAmGmCmAmAmUmAmGmUmC mCmCmAmUmAmG |
| 71 | 5'SCD55 | A*A*CCCCUACCAACUGGUCGGGGUUUGAAACGAUCACUGAGUCCUUCUCGCCAG |
| 72 | 3'SCD55 | AACCCCUACCAACUGGUCGGGGUUUGAAACGAUCACUGAGUCCUUCUCGCC*A*G |
| 73 | 5'-3'-SCD55 | A*A*CCCCUACCAACUGGUCGGGGUUUGAAACGAUCACUGAGUCCUUCUCGCC*A*G |
| 74 | 5'SeCD55 | U*U*UAACCCCUACCAACUGGUCGGGGUUUGAAACGAUCACUGAGUCCUUCUCGCCAG |
| 75 | 5'-3'-SeCD55 | U*U*UAACCCCUACCAACUGGUCGGGGUUUGAAACGAUCACUGAGUCCUUCUCGCCAGU*U*U |
| 76 | 5'S-3'SeCD55 | A*A*CCCCUACCAACUGGUCGGGGUUUGAAACGAUCACUGAGUCCUUCUCGCCAGU*U*U |
| 77 | 5'Se-3'SCD55 | U*U*UAACCCCUACCAACUGGUCGGGGUUUGAAACGAUCACUGAGUCCUUCUCGCC*A*G |
| 78 | 5'SCD71 | A*A*CCCCUACCAACUGGUCGGGGUUUGAAACCGAUCACAGCAAUAGUCCCAUAG |
| 79 | 3'SCD71 | AACCCCUACCAACUGGUCGGGGUUUGAAACCGAUCACAGCAAUAGUCCCAU*A*G |
| 80 | 5'-3'-SCD71 | A*A*CCCCUACCAACUGGUCGGGGUUUGAAACCGAUCACAGCAAUAGUCCCAU*A*G |
| 81 | 5'SeCD71 | U*U*UAACCCCUACCAACUGGUCGGGGUUUGAAACCGAUCACAGCAAUAGUCCCAUAG |
| 82 | 5'-3'-SeCD71 | U*U*UAACCCCUACCAACUGGUCGGGGUUUGAAACCGAUCACAGCAAUAGUCCCAUAGU*U*U |
| 83 | 5'S-3'SeCD71 | A*A*CCCCUACCAACUGGUCGGGGUUUGAAACCGAUCACAGCAAUAGUCCCAUAGU*U*U |
| 84 | 5'Se-3'SCD71 | U*U*UAACCCCUACCAACUGGUCGGGGUUUGAAACCGAUCACAGCAAUAGUCCCAU*A*G |
| 85 | iCD462 | AACCCCUACCAACUGGUCGGGGUUUGAAACAUAACAAUCACAGCAAUGACCCATd |
| 86 | 3'SCD462 | AACCCCUACCAACUGGUCGGGGUUUGAAACAUAACAAUCACAGCAAUGACC*C*A |
| 87 | iCD463 | AACCCCUACCAACUGGUCGGGGUUUGAAACUCACAAAUAGUAUGGGUGGCAAGTd |

| | | |
|---|---|---|
| 88 | CD463 | AACCCCUACCAACUGGUCGGGGUUUGAAACUCACAAAUAGUAUGGGUGGCA*A*G |
| 89 | iCD552 | AACCCCUACCAACUGGUCGGGGUUUGAAACACUCCACUGGACAGAGCUGCCUGTd |
| 90 | CD552 | AACCCCUACCAACUGGUCGGGGUUUGAAACACUCCACUGGACAGAGCUGCC*U*G |
| 91 | iCD553 | AACCCCUACCAACUGGUCGGGGUUUGAAACGCAAGCCCAUGGUUACUAGCGUCTd |
| 92 | 3'SCD553 | AACCCCUACCAACUGGUCGGGGUUUGAAACGCAAGCCCAUGGUUACUAGCG*U*C |
| 93 | iCD712 | AACCCCUACCAACUGGUCGGGGUUUGAAACUCCAUCAUUCUGAACUGCCACACTd |
| 94 | 3'SCD712 | AACCCCUACCAACUGGUCGGGGUUUGAAACUCCAUCAUUCUGAACUGCCAC*A*C |
| 95 | iCD713 | AACCCCUACCAACUGGUCGGGGUUUGAAACCAAGUUUCAAUAGGAGAGCUGUGTd |
| 96 | 3'SCD713 | AACCCCUACCAACUGGUCGGGGUUUGAAACCAAGUUUCAAUAGGAGAGCUG*U*G |
| 97 | 3'S-SCoV2NY1_g1 | AACCCCUACCAACUGGUCGGGGUUUGAAACAGAACAGAUCUACAAGAGAUC*G*A |
| 98 | 3'S-SCoV2NY1_g2 | AACCCCUACCAACUGGUCGGGGUUUGAAACAAGUUGGUUGGUUUGUUACCU*G*G |
| 99 | 3'S-SCoV2NY1_g3 | AACCCCUACCAACUGGUCGGGGUUUGAAACACAAGAGAUCGAAAGUUGGUU*G*G |
| 100 | LoopM-CD46 | AACCCCUACCmAmAmCmUGGUCGGGGUUUGAAACAGACAAUUGUGUCGCUGCCAUCG |
| 101 | LoopM-CD55 | AACCCCUACCmAmAmCmUGGUCGGGGUUUGAAACGAUCACUGAGUCCUUCUCGCCAG |
| 102 | LoopM-CD71 | AACCCCUACCmAmAmCmUGGUCGGGGUUUGAAACCGAUCACAGCAAUAGUCCCAUAG |
| 102 | LoopM-NT | AACCCCUACCmAmAmCmUGGUCGGGGUUUGAAACUACUUACCUAAUGCGCGGUAGAU |

<u>Modified bases</u>

m : 2'-O-methyl
* : phosphothioate bond
Td : inverted T

### Supplementary Table 2. PCR primers for T7 *in vitro* transcription.

| Primer | Sequence |
|---|---|
| CD46_IVT_F | *taatacgactcactataggg*AGTGTAAAGTGGTCAAATGTCGA |
| CD46_IVT_R | AAGACACTTTGGAACTGGGG |
| CD55_IVT_F | *taatacgactcactataggg*CTAATGCCCAGCCAGCTTTG |
| CD55_IVT_R | CAGCTACGATTGCAGAACTCT |
| CD71_IVT_F | *taatacgactcactataggg*TGTGGAGATGAAACTTGCTGT |
| CD71_IVT_R | ACCCCTTTACAATAGCCCAAGT |
| mRNA_RfxCas13d_F | *taatacgactcactataggg*GCCACCATGATCGAAAAAAAAAAGT |
| mRNA_RfxCas13d_R | CTAATTGCCGGACACCTTCTTTT |
| mRNA_eGFP_F | *taatacgactcactataggg*GCCACCATGGTGAGCAAGGGCGAGG |
| mRNA_eGFP_R | CTACTTGTACAGCTCGTCCATGC |

**Italicized nucleotides* indicate T7 primer binding site.

### Supplementary Table 3. Primers for SARS-CoV-2 TRS-Leader sequence.

| Primer | Sequence |
|---|---|
| Leader_F | CTAGCATTAAAGGTTTATACCTTCCCAGGTAACAAACCAACCAACTTTCGATCTCTTGTAGATCTGTTCTCTAAACGAACG |
| Leader_R | CTAGCGTTCGTTTAGAGAACAGATCTACAAGAGATCGAAAGTTGGTTGGTTTGTTACCTGGGAAGGTATAAACCTTTAATG |
